# Embryo timelapses can be compiled and quantified to understand canonical histone dynamics across multiple cell cycles

**DOI:** 10.1101/316281

**Authors:** Lydia Smith, Paul Maddox

## Abstract

In the last decade, computational analysis of big datasets has facilitated the processing of unprecedented quantities of collected biological data. Thus, automations and big data analyses have been revolutionary in detecting and quantifying subtle phenotypes in cell biological contexts. Analyzing similar quantities of data in larger and more complicated biological systems such as live embryos has been more challenging due to experimental necessities impeding both compilations of data collection and informative analysis. Here we present a streamlined workflow that can quantify cell cycle dynamics in early developing embryos using fluorescently labeled proteins. We benchmark this pipeline using *Caenorhabditis elegans* (nematode) embryonic development and a fluorescently labeled histone. Using our pipeline, we find that histone proteins are broadly stable in early embryonic development. In sum, we have utilized the large biological and experimental variation associated with quantification of fluorescent proteins in embryonic systems, to quantify nuclear accumulation rate, chromatin incorporation, and turnover/stability of canonical histones during early development.

## Introduction

Measurement of the quantities and dynamics of chromatin organizing proteins throughout the cell cycle is a widely used method in cell biology. Biochemical measurements of cell cycle components in yeast or mammalian cultured cells rely on immunoblots of synchronized cultures of hundreds or thousands of cells (Foltz et al. 2009; Juanes 2017). More recently, high resolution and magnification (NA >1, 60x^+^ magnification) fluorescence microscopy has been utilized as an alternative method of measuring levels of fluorescently labeled proteins during the cell cycle. One of the advantages of fluorescence microscopy is it allows for the quantification of proteins in living cells, using either fluorescently labeled proteins or active dyes at the single cell level. Mammalian cell cultures and yeast have been the workhorses of *in vivo* quantitative fluorescent microscopy, due to the plethora of molecular tools available enabling fluorescent tagging of proteins, insertion of expression vectors, and the introduction of active dyes into cells. They are also ideal for imaging as they have little movement in 3D, have relatively thin vertical profiles, and fluorescent proteins are in close proximity of the coverslip, resulting in a high signal to noise ratio (Mattheyses et al. 2010; Jansen et al. 2007). Cultured cells exhibit synchronous and symmetric cell cycles making statistical analysis of collected data easy to perform. Due in part to the highly stereotyped nature of these systems, much of the data collection process can now be automated (Roukos et al. 2015; Wang et al. 2017).

In order to measure histone dynamics in eukaryotic embryos, it is essential to accurately measure cell cycle progression. Quantitative fluorescent microscopy presents challenges compared to cell cultures, including non-uniform cell architecture (variable sizes and shapes), typically short, rapid, often asynchronous, cell cycles (followed by increasingly lengthening cell cycles (Philpott & Yew 2005)), an inability to pause or synchronize cell cycles (due to weak cell cycle checkpoints (Kipreos 2005) or impermeable eggshells (Huang et al. 2015)), and significant movement of cells/nuclei throughout development (Ishiura 2010).

*C. elegans* was first developed as a model system by Sydney Brenner and has become a workhorse for embryology studies due to its transparent egg shell and invariant development (Byerly et al. 1976; Sulston et al. 1983). In addition, *C. elegans* was the first eukaryotic organism in which fluorescent protein GFP was visualized (Chalfie et al. 1994) ushering in the capability for automated tracking of fluorescent proteins throughout development (Boyle et al. 2006). Like many other embryonic systems used in the laboratory (Xenopoulos et al. 2012), *C. elegans* embryos have several physical characteristics that make quantification of fluorescence microscopy experiments challenging to perform. First, whereas the phases of mitosis are easy to temporally distinguish due to the condensation and behavior of chromosomes, *C. elegans* embryos lack conventional gap phases, making it difficult to visualize unique events in between. Second, it is virtually impossible to collect synchronized populations of cells/embryos, precluding bulk biochemical and/or immunofluorescence analyses. Third, *C. elegans* embryos are ~50um by ~20-25μm ellipsoids through which cells and nuclei may migrate, either passively through progressive cell cycles, or actively through tissue migration events like gastrulation (Byerly et al. 1976; Young et al. 1991; Schnabel et al. 2006). Thus, the fluorophore signal throughout the embryo decreases exponentially deeper into the sample when using the required high Numerical Aperture (NA>1.0) optics (Waters 2009).

When using quantification of image data-sets generated using light microscopy to measure protein levels, there are inherently two large sources of variation within the data: biological and experimental. Biological variability can be minimized by maintaining consistent imaging and culturing conditions (Byerly et al. 1976), but it is an aspect of the data that is inherent to biological specimens Therefore, we focused our analysis efforts on eliminating experimental variation in our analysis in order to create a model of protein dynamics in an “average embryo”. In cell culture, where a relatively homogenous group of cells can be synchronized, it is possible to easily average quantifiable information about the cells at specific parts of the cell cycle. It is also possible to sort non-synchronized cells either through similarity in cell cycle progression (every cell progresses through the cell cycle identically allowing post-acquisition alignment of data (Gookin et al. 2017)) or bulk snap shot analysis. Here we have exploited the stereotypical development in *C. elegans* to create an analysis pipeline to quantify histone dynamics through post-acquisition alignment of data. The *C. elegans* system allows the comparison of multiple differentiated lineages from as early as the 2-cell stage until hatching and has made it possible to analyze embryogenesis using either differential interference contrast (DIC) microscopy or fluorescently labeled proteins (Boyle et al. 2006). This ease of visibility of all of the cells of the developing embryo has been instrumental in developing our understanding of the spatial and temporal regulation of many proteins, transcripts, and developmental factors.

To perform post-acquisition alignment of data, we took advantage of *C. elegans’* stereotypical development by utilizing embryonic equivalence. We define embryonic equivalence as the assumption that in highly stereotypical embryogenesis such as *C. elegans* (Sulston et al. 1983; Boyle et al. 2006), developmentally identical cells between embryos will have equivalent biological measurables; this includes cell cycle length, size, position, protein levels, etc. This embryonic equivalence allows us to average measurements from multiple embryos in order to reduce experimental variability and build a model of an average embryo. As long as all the timelines share at least one common cellular event, all time-lapses can be both normalized and organized temporally together. Because the length of the cell cycles fluctuates a small amount based on environmental and imaging conditions, we aligned the timelapses to the last time point of a cell’s lifespan. In our case, this was determined by the last timepoint when the mitotic chromatin was still one mass, and not divided into two groups during anaphase (Figure 1A). This approach has allowed us to quantify histones in early development and could easily be transformed to other cellular components (e.g. microtubules, Golgi, ER, etc.) in *C. elegans* or other complex cellular systems such as embryos or organoids.

**Figure 1.**
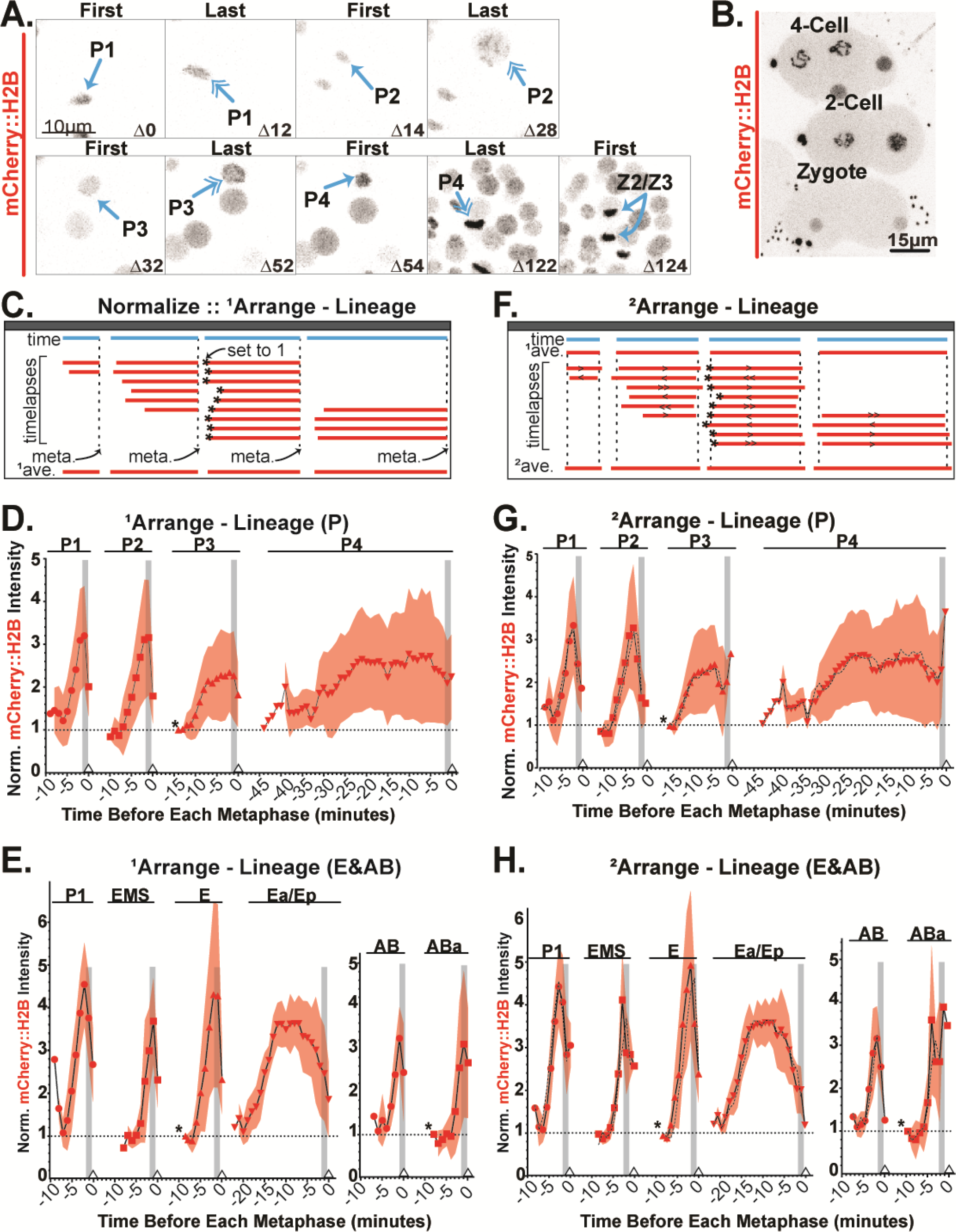
Embryo analysis pipeline involving normalization, organization, and secondary analysis facilitate quantification of canonical histone behavior in C. elegans embryos. (A) Representative image of a timelapse image of the 1-cell through ~100-cell embryo. The P-lineage is indicated and representative first and lapse timepoints for each cell cycle are shown. Developmental time indicated in minutes post 2-cell stage. (B) Representative image of multiple, asynchronous embryos imaged simultaneously. (C) Schematic of data normalization and primary arrangements. For lineage arrangements, all cell cycles are aligned to the last timepoint regardless of cell cycle length (~metaphase = meta.). All timelapses must contain a shared cell cycle event (*), the nuclear value at this point was set to 1 within each timelapse. Resulting primary ^1^averages can be then be calculated. (D-E) Timepoints are 2-minutes apart. Grey column represents approximate time of NEBD. Small triangle indicates alignment point. Experimental ^1^averages and standard deviations are plotted. (D) P-lineage experimental data plotted. (*) is the first timepoint of P3. (E) E-lineage (left) and AB-lineage (right) experimental data plotted. (*) is first timepoint of E and ABa respectively. (F) Using the calculated ^1^average and the same data sets, the data is run through our custom MATLAB macro to generate new timestamps to decrease the standard deviation around the ^1^average, resulting in a new ^2^average generated as a histogram with 1 timepoint (2-minute) bins. (G-H) Experimental ^1^averages (black dashed line), ^2^averages, and standard deviations are plotted. (G) P-lineage experimental data plotted. (H) E-lineage (left) and AB-lineage (right) experimental data plotted.

## Reagents and Instruments

### C. elegans strains

Worms were cultured and imaged at 20°^c^ on OP50 seeded agar plates as described in Riddle *et al* (Riddle et al. 1997). All worms used for imaging were from non-starved populations. The *C. elegans* strains used here (OD421) were a generous gift from the Desai and Oegema labs. OD421 is a triple mutant composed of a composition of the following strains: CeCENP-*A/hcp-3* deletion allele *ok1892*, GFP-CeCENP-A (OD347), and *pie-1p*:mCherry-histone H2B (OD56)(Gassmann et al. 2012).

### Microscope

Imaging was performed on a Nikon A1R microscope body with a 60X 1.27 NA Nikon Water Immersion Objective with a GaASP PMT detector (Nikon) using NIS-elements.

## Methods

### Microscopy

#### Cell Identification

Cell identification was performed through manual lineage tracing from at least the 4-cell stage in the early *C. elegans* embryos. The late P-lineage identification was aided by the extra fluorescence in the red channel driven by the pie-1 promoter driving mCherry::H2B. Cellular morphology, movement, and cell cycle duration was used to confirm the identification of cells.

#### Sample preparation and imaging of embryos

Our protocol is adapted from Monica Driscoll’s protocol on Wormbook. Embryos were dissected out of non-starved, gravid adult *C. elegans* into M9 on No.1.5 22mm^2^ coverslips and mounted onto 2% (w/v) agar pads on a standard microscopy slide, before being sealed with VALAP (1:1:1 lanolin, petroleum jelly, and parafilm wax). 2% agar pads were used to gently compress embryos without damaging them. Imaging was done using excitation of mCherry fluorophores using a 561nm laser. Z-stacks contained between 20-30 slices (depending on how embryo was oriented), and all slices were 1μm apart. Z-stacks were collected every 2 minutes for approximately 1-3 hours for early embryo timelapses, and for the entire duration of each cell cycle for FRAP analysis. Images collected using NIS-Elements were imported for analysis in ImageJ/FIJI (Schindelin et al. 2012).

## Results

### Embryonic time-lapse data can be organized to create a model of an “average” embryo

To quantify the dynamics and quantities of histones in developing embryos, we utilized a strain of *C. elegans* that has one copy of H2B *(his-58)* tagged with mCherry. Each cell cycle, the amount of H2B in each nucleus or incorporated into chromatin is measured over time by fluorescence intensity. We focused primarily on three early lineages; two somatic lineages, AB and E, as well as the germline lineage P (Figure 1A). Embryos were imaged within 3D 1.0μm interval Z-stacks with 2-minute acquisition intervals for 1-3 hours. Multiple non-synchronized embryos were imaged simultaneously to maximize data gathered per image (Figure 1B). All Z-stacks were transformed into individual Max-Projections throughout the entire embryo before manually drawing regions of interest (ROIs) and taking measurements using FIJI/ImageJ (Schindelin et al. 2012). If a Max-Intensity-Projection throughout the entire embryo was not appropriate for measuring a single nucleus in a crowded embryo, individual Max-Projections encompassing only that nucleus were generated for each time point. For every timepoint, a ROI was drawn around the entire nucleus of interest, and two or three ROIs of equal size were collected from the cytoplasm. Collection of measurements were expedited using custom macros. Each nucleus and cytoplasm (background) of the appropriate lineages from every embryo is quantified, background and bleach corrected (Equations 1–2) and given a timestamp relative to anaphase.

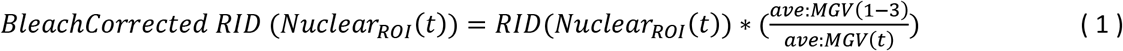

This equation generates the RawIntDen (RID) of a nucleus of interest calculated with a hand-drawn ellipses Regions of Interest (ROI). To bleach correct this value over the course of a timelapse, the RID of the nucleus is multiplied by the ratio of the Mean Grey Value (MGV) of the background ROIs by the average MGV of the first three timepoints. This was to account for any variability in the background signal of the first three timepoints. This bleach correction term becomes larger in much later timepoints as more bleaching occurs.

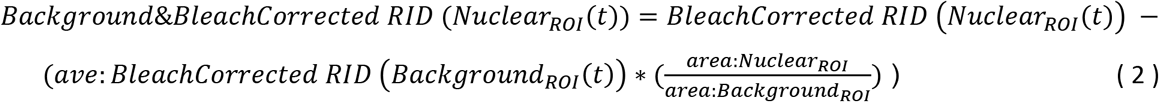

This equation generates the RIDs that are used in Figures 1–2. Background ROIs are multiplied by the same MGV ratio term used in Equation 1 to correct for bleaching during the timelapse. The background RID term does have an additional modifier that accounts for the smaller size of the background ROIs compared to the nuclear ROI. Because the background ROIs must sometimes be smaller than the nuclear ROI, especially in smaller cells, the background RID must be scaled up appropriately before being subtracted from the nuclear RID in Equation 2. Example image in Supplemental Figure 1.

For our primary analysis, all timestamps are aligned to the last time-point before each cell’s anaphase (i.e. approximately metaphase) with the last image being set to 0. The calculated time-lapse RIDs can also be normalized further on a lineage specific basis. When normalized on a lineage by lineage basis, the values at a shared cellular event are set to 1, and all other values in that lineage are normalized accordingly (Figure 1C). From either of these analyses an average value can be generated at each time-point to build a profile of the relative quantities of a histone throughout cell cycles and lineages. Experimental data from the germline P-lineage (Figure 1D) and the somatic E- and AB-lineages (Figure 1E) are shown.

In order to image the embryos for long periods without significantly bleaching or inducing phototoxicity within the embryos, we acquired stacks of images through the embryos every two minutes. To overcome possible temporal under-sampling of the data, we developed a MATLAB macro that automatically explores all the possible combinations of aligning the timelapses in an attempt to decrease the overall standard deviation around the average we had previously generated. This macro allows us to interleave the timelapses by fitting similarly shaped curves closer mathematically (Berro & Pollard 2014; Boudreau et al. 2018 Preprint). All the timelapses maintain their normalized values, and time-points are maintained at 2 minutes apart. However, the timelapses of each cell cycle are shifted within a 4-minute (±2 minutes) interval by the algorithm either forward (>, >>) or backwards (<, <<) in time by either a small (>, <) or large amount (>>, <<) (Figure 1F). Actual experimental data for the P-lineage are shown in Figure 1G, and somatic E- and AB-lineages in Figure 1H.

To determine the dynamics of H2B in each cell cycle, we compared the relative last timepoint values of H2B signal derived from our “average embryo” pipeline (Figure 2A). To calculate the rate of histone nuclear accumulation during all of the observed cell cycles, we quantified the duration of each cell cycle wherein the nuclear signal rose in minutes (Table 1 Columns 1-2). We then normalized all of the timelapses to the relative value at each lineage’s representative at the 4-cell stage (Table 1 Column 3). This number was multiplied by the amount of histone to be added during DNA replication and then divided by the import duration to calculate nuclear accumulation (Table 1 Columns 4-5 and Equation 3). Import durations are manually measured for each cell cycle encompassing when the signal first begins to rise to when it stabilizes or the cell cycle ends.

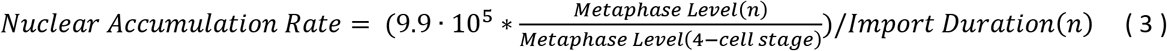

**Figure 2.**
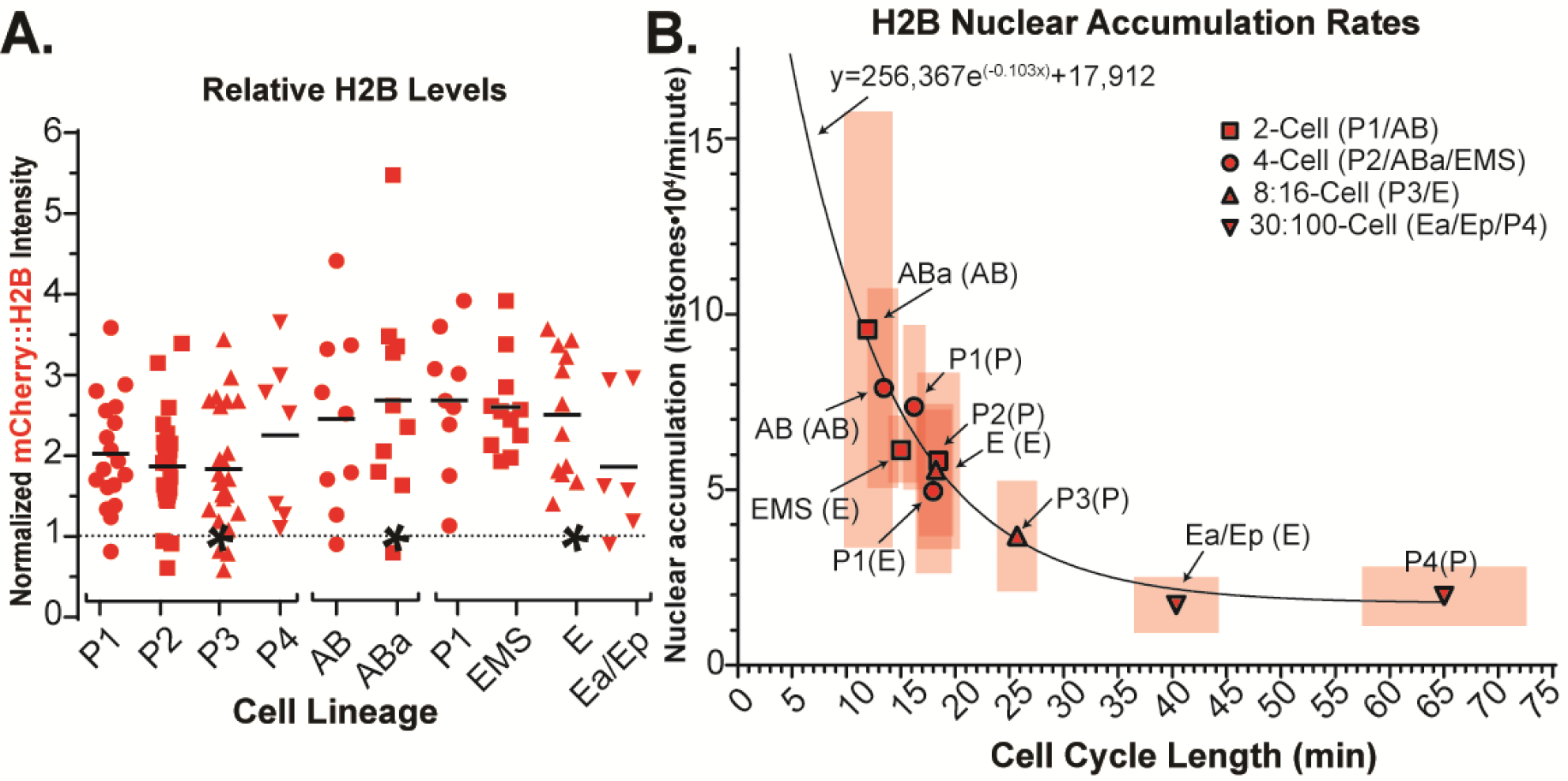
Histone nuclear import rates scale inversely to cell cycle length. (A) A germline-specific-promoter drives increased fluorescently labeled histone in the P-lineage, whereas the amount of fluorescently labeled histone incorporated into chromatin is stable through the 4-cell stage and decreases starting around the 8-cell stage of the somatic lineages. Identical data to Figure 1D-E each normalized internally to their own lineages (*). (B) Plotted calculations of H2B nuclear accumulation rates throughout development. Boxes represent standard deviation of the mean for both cell cycle length (horizontal) and nuclear accumulation rate (vertical). Solid line is a best fit one phase exponential decay curve, equation shown.

**Table 1.** List of calculated values for quantifying nuclear accumulation of H2B. (1) Cell cycle analyzed. (2) Rise in nuclear signal measured for each cell cycle of the three lineages quantified in minutes. (3) Relative levels of mCherry::H2B from ^1^lineage analysis (Figure 1) (left), then normalized to the cell value from the 4-cell embryo (P2/EMS/ABa) (right). (4) Calculated number of histones with 1.00 = 9.9•10^5^ histones. (5) Values in Column 3 and multiplied by values in Column 4. (6) Number of timelapses used to generate the calculated accumulation.

| <sup>1</sup>Cell | <sup>2</sup>Import Duration | <sup>3</sup>Relative Metaphase Levels (versus 4-cell) |  | <sup>4</sup>Histone Added | <sup>5</sup>Nuclear Accumulation (histones/min) | <sup>6</sup>n |
| --- | --- | --- | --- | --- | --- | --- |
| P1 | 12.3 (± 2.13) | 2.02 | 1.09 | 1.08•10<sup>6</sup> | 7.36 (± 2.27)•10<sup>4</sup> | 10 |
| P2 | 13.6 (± 1.79) | <u>1.86</u> | 1.00 | <u>0.99•10<sup>6</sup></u> | 6.08 (± 2.52)•10<sup>4</sup> | 18 |
| P3 | 12.6 (± 3.54) | 1.83 | 0.98 | 0.97•10<sup>6</sup> | 3.74 (± 1.28)•10<sup>4</sup> | 20 |
| P4 | 15.6 (± 5.17) | 2.16 | 1.16 | 1.15•10<sup>6</sup> | 1.79 (± 0.76)•10<sup>4</sup> | 6 |
| P1 | 11 (± 2.58) | 2.68 | 1.03 | 1.02•10<sup>6</sup> | 4.98 (± 2.34)•10<sup>4</sup> | 4 |
| EMS | 9.8 (± 2.52) | <u>2.59</u> | 1.00 | <u>0.99•10<sup>6</sup></u> | 6.19 (± 1.03)•10<sup>4</sup> | 10 |
| E | 12.0 (± 3.13) | 2.50 | 0.96 | 0.95•10<sup>6</sup> | 5.33 (± 1.96)•10<sup>4</sup> | 9 |
| Ea/Ep | 12.3 (± 1.96) | 1.86 | 0.72 | 0.71•10<sup>6</sup> | 1.75 (± 0.80)•10<sup>4</sup> | 6 |
| AB | 6.8 (± 1.51) | 2.45 | 0.91 | 0.91•10<sup>6</sup> | 7.94 (± 2.85)•10<sup>4</sup> | 7 |
| ABa | 7.7 (± 1.56) | <u>2.68</u> | 1.00 | <u>0.99•10<sup>6</sup></u> | 9.61 (± 6.22)•10<sup>4</sup> | 9 |

For this calculation we made the assumption that the histones roughly double every cell cycle and that there are roughly 1 million H2B histones added each cell cycle. This is based on *C. elegans’* 100Mb diploid genome size (Hillier et al. 2005) divided by 147bp wrapped around each histone with an average 50bp DNA linker (Szerlong & Hansen 2011) with two H2B histones per nucleosome. Interestingly, we found an exponential decrease of nuclear accumulation rates as cell cycle length increases; a 6.7 min decrease half-life of accumulation for our H2B nuclear accumulation curve indicating that with every 6.7-minute increase to a cell cycle’s length, there is a halving of the rate of nuclear accumulation (Figure 2B).

### *C. elegans* canonical histones are stably incorporated during early embryogenesis

Given that the behaviors of histones in *C. elegans* embryos partially differed from canonical systems (humans, yeast, frog etc), to further probe these behaviors we turned to a different fluorescent assay, Fluorescence Redistribution After Photobleaching (FRAP)(Walczak et al. 2010). Utilizing the same strain of *C. elegans* described above, we focused our efforts on measuring the turnover of histones from the beginning to end of each cell cycle in early embryos. The most common method of measuring protein stability and turnover rates, FRAP, involves photobleaching all of the fluorophores in a particular region of interest and measuring both the quantity and rate of recovered fluorescent signal in that area (Supplemental Figure 2A). Typically, these types of experiments involve an internal non-bleached control. In our case, we wanted to bleach the entire pool of fluorophores in the nucleus, meaning that there would be no corresponding non-bleached homologous region to compare it to. Instead of traditional controls, we used the nuclear signal of non-bleached embryos as our control (Figure 3A).

**Figure 3.**
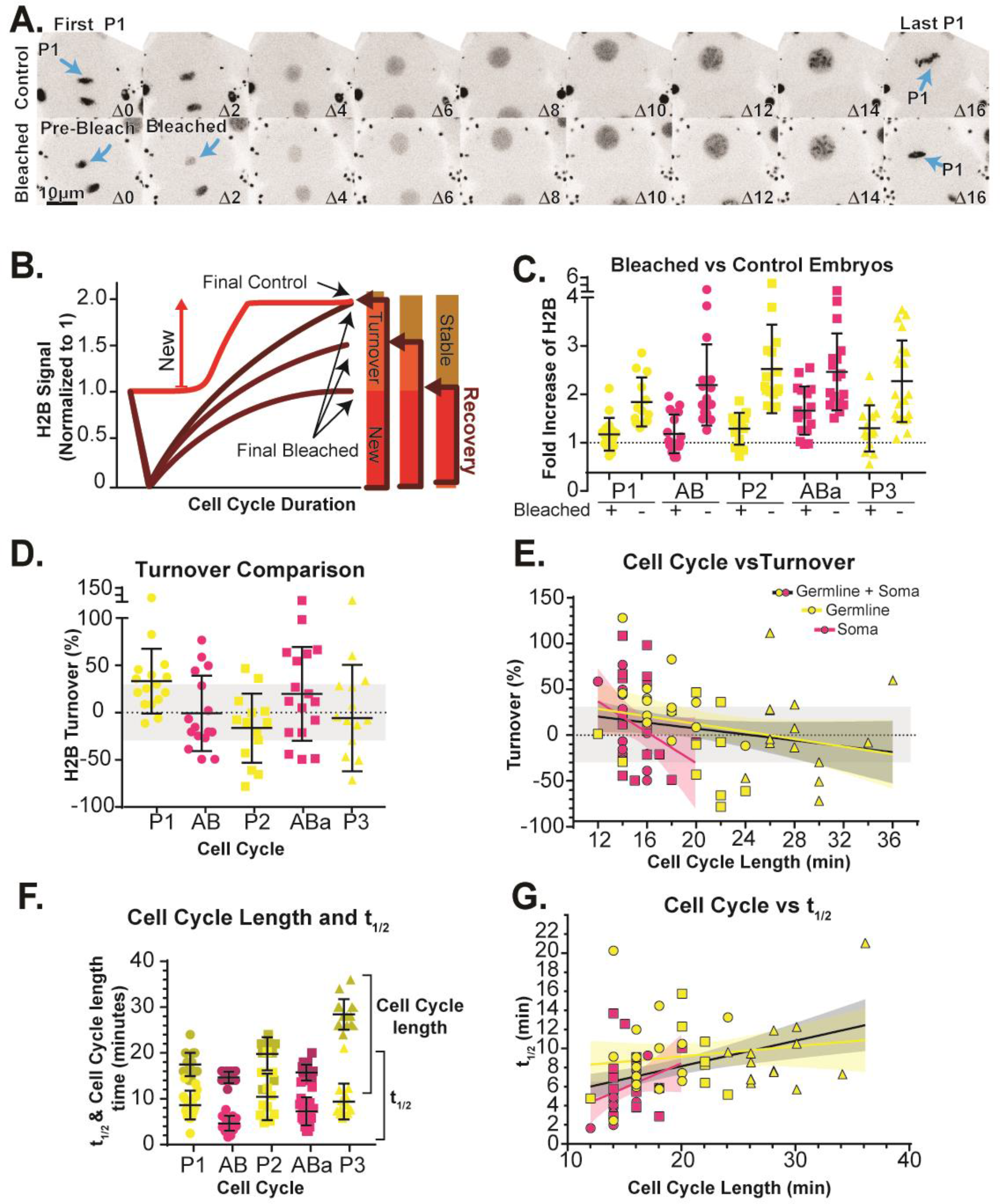
Protein stability and turnover dynamics can be quantified using population controls to reveal stable chromatin incorporation. (A) Max-projections of representative images of Control (top) and Bleached (bottom) embryos. Images are scaled and aligned identically with first and last timeframes of specific cells (P1) indicated. (B) Example of where in each timelapse data is normalized and collected from to calculate turnover. (C) Experimental values of Final Values for Control and Bleached conditions for 5 different cells. (D) Calculated turnover values for 5 cells from early embryo. (E-G) Linear regression lines for somatic cells (pink), germline cells (yellow), and both (black) with 95% confidence interval bands. (E) Turnover values plotted against cell cycle length. (F) Calculated values of Cell Cycle length and t_1/2_ (G) Cell cycle length plotted against calculated W2.

To determine H2B turnover and accumulation in the nucleus, the amount of fluorescent signal in the nucleus was measured over the cell cycle in bleached and unbleached samples. In order to calculate the fraction of recovered signal, we normalized all of the timelapses to their first time-point (no bleach correcting) and calculated the increase in the controls (Figure 3B-C and Equation 4), then subtracted that average increase from each of our recovery FRAP signal turnover (Figure 3D).

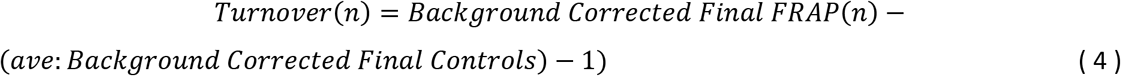

This final value was calculated and only values with a FRAP efficiency of at least 70% were kept (meaning at least 70% of the measured signal disappeared post-laser ablation with at most 30% remaining). We find that all of the cells of the early embryo examined fall within the efficiency margin around 0%. We also find that this characteristic does not change when we separate the cells based on cell cycle length/developmental stage instead of cell cycle (Figure 3E). From the FRAP curves, we can also calculate the rate at which the signal recovers in each cell cycle (Maddox et al. 2000) (Supplemental Figure 2B-C) as well as the inverse of how long it takes each cell to recover their signal, t_1/2_ (Figure 3F-G and Equations 5–6. RID = RawIntDen).

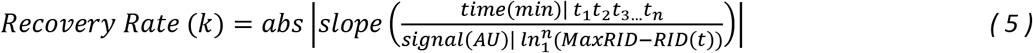

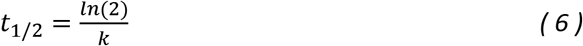

For either measurement, there is not a strong correlation between cell type (soma/germ), cell cycle length, or developmental stage and the very slow rate at which the cells replace their old histones (Figure 3E and G).

## Discussion

Here we used a quantitative microscopy analysis pipeline to investigate histone dynamics in early embryonic development. As genome activity changes, so does histone stability; transcriptionally silent regions have more histone stability (Kireeva et al. 2002). In *C. elegans* embryos, transcription is activated in varying lineages at differing times (Tintori et al. 2016), thus it could be possible to detect these changes by probing histone population dynamics. We have found that gross histone dynamics are canonical (follow that reported in other model systems) in *C. elegans* with very little variation in differing lineages. This result could indicate that genome regulation in early development leads to histone stabilization, or that these changes are highly focused and thus not detected in our assays. Despite this, our work has derived a new imaging analysis pipeline and shows that histones in rapidly dividing embryonic cells generally follow expected behaviors.

One persistent complication to studying protein dynamics in embryos using light microscopy is the large amount of experimental variability that is added into any quantitative analysis attempted. Because the cells in a developing embryo are more demanding in their preferred environmental conditions than traditional cell cultures, creating an ideal quantitative environment has been a challenge for quantitative microscopists. In order to generate an ‘average embryo’, we found that normalizing, aligning, and averaging of many individual embryo timelapses is an effective way to understand the dynamics of cell cycle proteins throughout early embryo cell cycles. The resulting analyses can be differentially organized to answer specific biological questions. In our case, we were able to create models of ‘average embryos’ in early *C. elegans* embryos and describe the protein dynamics of individual cells/lineages. Utilizing our quantification and analysis pipeline, we focused our efforts on determining how H2B levels fluctuate throughout the cell cycles of developing lineages and if their characteristics were conserved in the early embryo. We chose H2B as our initial protein to start with given that it is an incredibly well conserved protein across phylogeny (Malik & Henikoff 2003) with well characterized cell cycle dynamics and characteristics (Osley 1991).

Although our fluorescent histone is driven by a germline specific promoter (pie-1), it is unclear exactly why such a strong and consistent amount of signal is present in all of the somatic cells (Fukuyama et al. 2006). It is possible that the promoter itself is ‘leaky’, meaning there is transcription of the gene in cells where we wouldn’t predict any to be. This could explain why in the germline (P) lineage analysis there is a slight increase in the amount of fluorescent histone incorporated into the chromatin at metaphase in the later cells, and this is not seen in the other somatic (AB and E) lineages (Figure 1D-E and Figure 2A). This mechanism would only explain changes we see in nuclear levels of the histones of later, post-30-cell, embryos as transcription does not become activated until after this point (Robertson & Lin 2015; Spencer et al. 2011). The more likely explanation in the early embryo is there is a large amount of maternally loaded mRNA in the zygote that is utilized by the embryo. Because of this, we think that for each cell cycle a certain amount of the pre-loaded mRNA is translated and imported into the nucleus and that the amount of fluorescent signal generated each cell cycle is not enough to measurably change our background signal given how large the volume of the cytoplasm is, especially in the early cell cycles.

This promoter could also explain why we see an ‘overloading’ of fluorescent histone into the nucleus, which then dissipates into the cytoplasm upon NEBD. This is why we predict that we see a stereotypical drop in intensity every cell cycle right before each metaphase. These ‘extra’ histones are not deleterious to mitotic fidelity as we see no impact on hatch/growth rate or incidence of males in this strain of worms (data not shown)(Hodgkin et al. 1979). We also note that even though the levels of the histone are different between the different lineages; although, each lineage is normalized to itself, allowing us to correct for these differences. Despite these potential limitations to our system, we find that the dynamics and characteristics of H2B in this system behave as we would predict them to.

We confirmed that throughout all of the cell cycles analyzed in early development, there was the expected rise of nuclear levels within each cycle cell from beginning to end. This rise in nuclear levels of canonical histones is always associated with S-Phase, when histones are transcribed, translated, and imported into the nucleus for incorporation into newly replicated chromatin. In *C. elegans* we see the rise occurring in the very early part of each cell cycle, usually occurring within the first ten minutes of the beginning of each cell cycle. Given that all of the early cells in the embryo lack GAP phases, DNA replication takes the entirety of the interphase cell cycle, and this is consistent with the accumulation of nuclear signal soon after mitosis terminates.

We also find that the data alignment application results in smoother curves (when n is high enough) that are potentially more physiologically accurate than data aligned strictly by their timepoints. However, the somatic cell cycles, which were composed of smaller datasets, had averages that seemed more susceptible to outliers, especially near the beginning and ends of the timelapses. This is because as the program explores all the possible ways to shift the timepoints of the datasets, it might reduce the standard deviation around the mean, but may cause points at the beginning or end of the timelapses to deviate from the average.

FRAP analysis showed that using unbleached embryos as a control group is sufficient to calculate turnover in our system. We find that although there is variation in the quantity of fluorescent histones measured per cell, overall the average in the controls is double the starting value, which is consistent with our understanding of canonical histone incorporation. Unless there is a significant change to the genome size, there will not be a change in the quantity of histones that are required each cell cycle. Because the controls were all imaged identically, they were averaged together to create a Final (Control) value. Because each photobleached embryo was bleached to a different percent of its original signal (due to movement of the nucleus during the bleaching process) each time lapse was one Final (FRAP) value. From these calculations and the fact that our average bleach efficiency of the original signal was around 70%, we found that all of the turnovers fell within that error margin. So, although there may be some amount of turnover, it’s not obviously apparent exactly how much that is, although we definitely do not have a complete recovery of signal as would be expected for a turnover of 100%. The stability of the histones means that the short and relatively unchanging recovery rates are a reflection of how long the turnover takes from the beginning after the cell cycle starts, and the time it takes to completely turnover does lengthen slightly with the lengthening of the cell cycle. With the exception of a few outliers, there really isn’t any strong correlation in any of the plots, suggesting that turnover is quite stable during the first few cell divisions and between lineages. We suspect that the histones are relatively stable in these early cell divisions because there is little to no detectable transcription occurring during any of the cell cycles that we performed FRAP experiments on. We predict, that if we were able to perform the FRAP experiments in the much longer somatic lineages (P-lineage has no transcription(Ghosh & Seydoux 2008)) that we might see some measurable turnover of the histones.

Overall, we find that utilizing an extremely well-studied protein in a highly characterized model system has been ideal for creating an embryonic experimental system where we can combine techniques from a variety of established cell cycle analysis pipelines for exploring protein dynamics and turnover. The analysis pipeline we have described here can be applied to any fluorescently labeled protein with a reasonable signal to noise ratio. It can be utilized in systems were temporal and spatial resolution are not cutting edge, or in systems where traditional synchronization and bulk biochemical analyses have been unable to be performed before. Although there is a larger amount of variation within the data than is usually appreciated in these types of experiments, information about the biology of these systems can still be produced, interpreted and appreciated despite their ‘setbacks’.

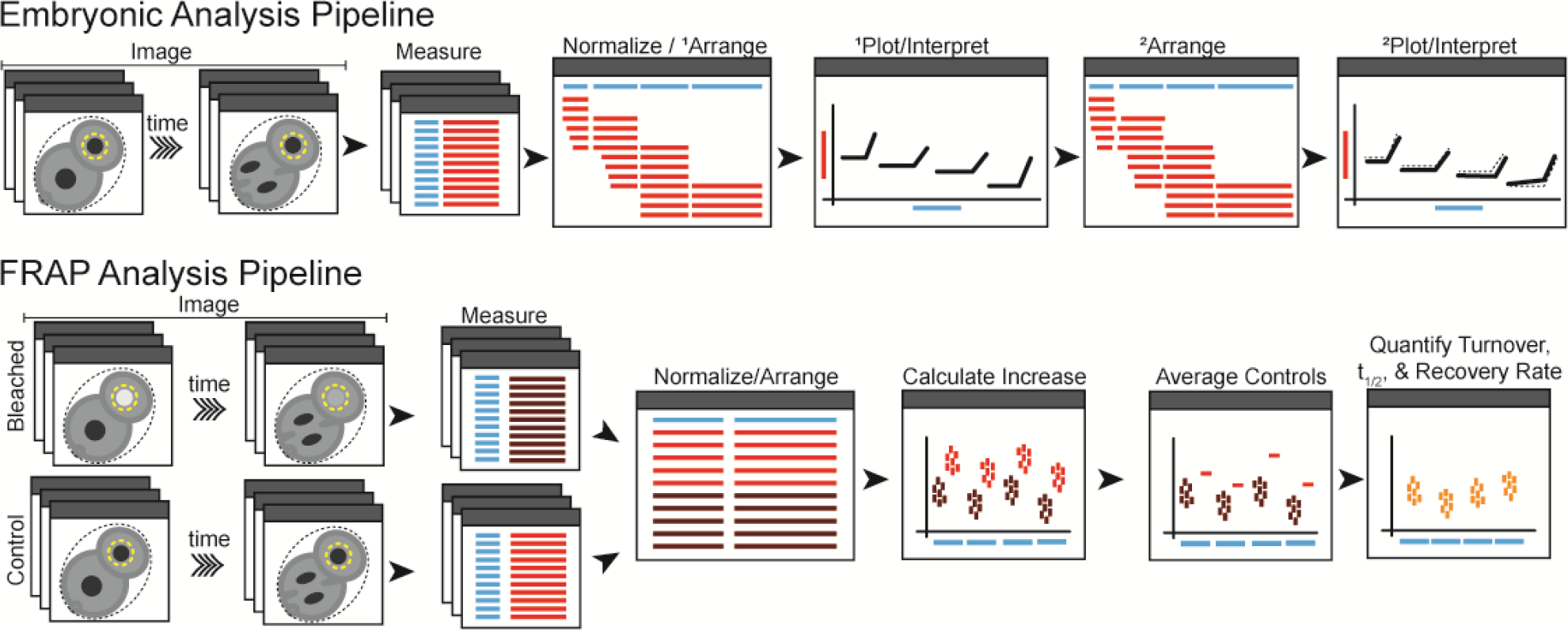

## Supplemental Material Legends

**Supplemental Figure 1.**
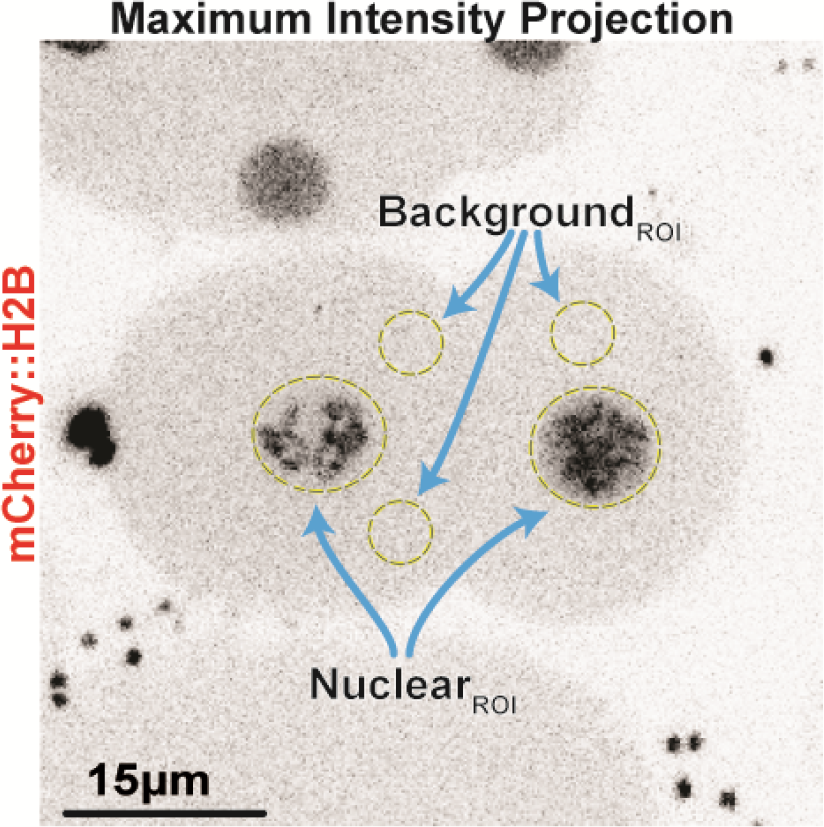
Representative image of manually collecting ROIs for measurement/quantification. NuclearROIs refer to drawing an ROI around a circular nucleus (pre-NEBD) (left) or chromatin (post-NEBD) (right). Background_ROI_s refer to drawing an ROI anywhere in the cytoplasm of the cell/embryo being analyzed.

**Supplemental Figure 2.**
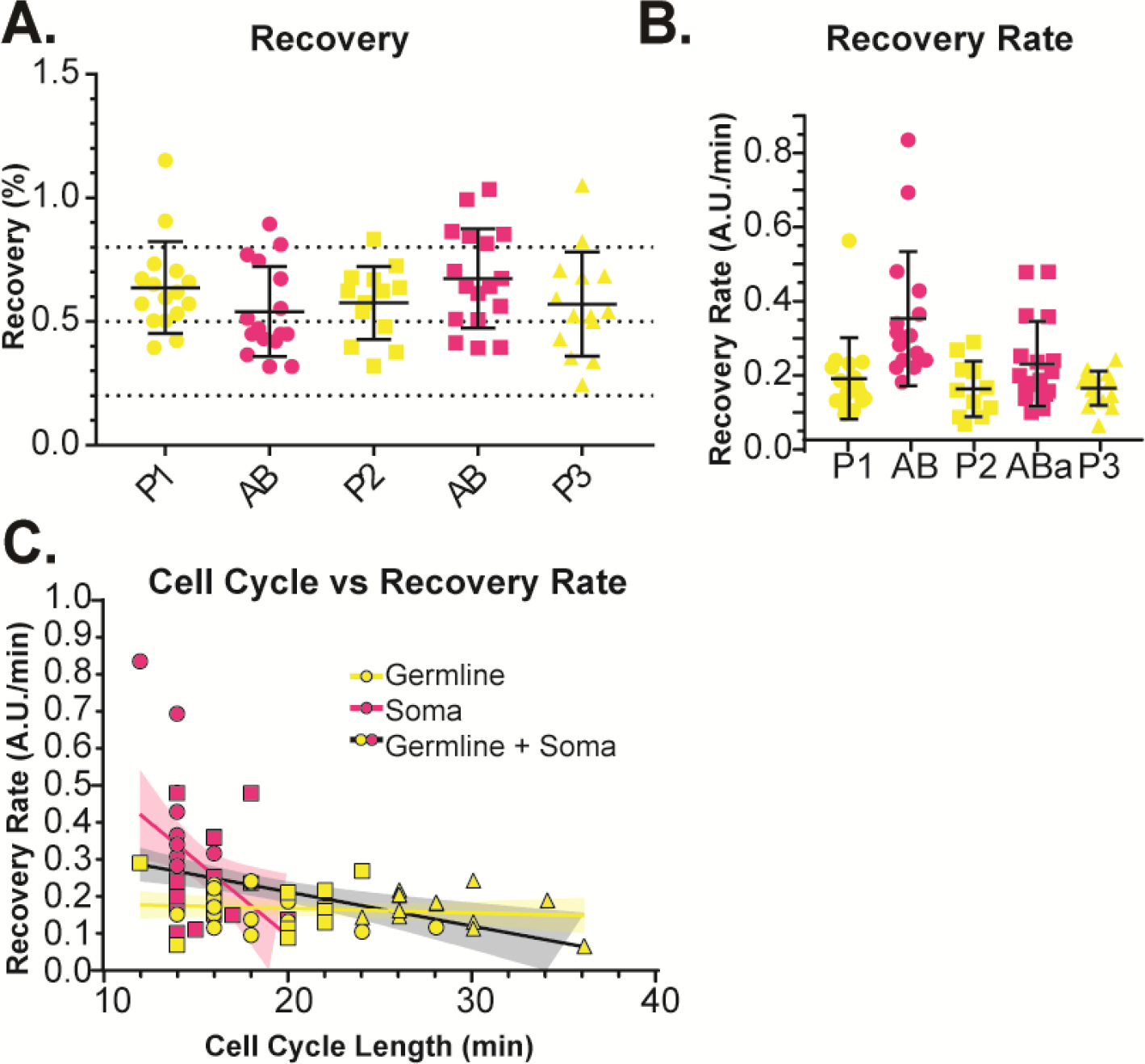
Protein recovery and recovery rate calculations. (A) Calculated recovery of signal in bleached embryos using only the first and final timepoints to calculate recovery of signal. (B) Calculated recovery rates using Equation 5. (C) Cell cycle length versus calculated recovery rate. Linear regression lines for somatic cells (pink), germline cells (yellow), and both (black) with 95% confidence interval bands.

## Acknowledgements

We would like to give a huge thanks to all the members of both Paul Maddox and Amy Maddox’s labs especially to Vincent Boudreau and Jake Sellinger who read this manuscript and helped develop the MATLAB macro used respectively. Special thanks to Kerry Bloom and Steve Rogers for their mentorship and guidance as well as reading this manuscript. We would also like to thank the Oegema/Desai labs for the worm strain used in this publication. This work was funded by the National Science Foundation CAREER Award 1652512.

## Author Contributions

*Conceived and designed the experiments:* Lydia Smith and Paul Maddox. *Performed the experiments:* Lydia Smith. *Analyzed the data:* Lydia Smith. *Wrote the paper:* Lydia Smith and Paul Maddox.

## Abbreviations Used

AU: Arbitrary Units
Ave: Average
FRAP: Fluorescence Recovery After Photobleaching
H2B: Histone 2B
Max-Projection: Maximum-Intensity-Projection
MGV: Mean Grey Value
NEBD: Nuclear Envelope BreakDown
RID: RawIntDen: Raw Integrated Density
ROI: Region of Interest

